# Volitional stopping is preceded by a transient beta oscillation

**DOI:** 10.1101/2024.12.28.630599

**Authors:** Joana F. Doutel Figueira, Reetta A. Ojala, Dmitrii Vasilev, Alejandro De Miguel, Lucas Jeay-Bizot, Ryo Iwai, Isabel Raposo, Negar Safaei, Lilian de Sardenberg Schmid, Uri Maoz, Masataka Watanabe, Nelson K. Totah

**Author notes:** Corresponding author: Nelson Totah. JFDF and RAO are equal contributors and listed in alphabetical order.

## Abstract

Human motor cortex EEG beta (15-30 Hz) oscillations undergo transient power modulations (bursts) during volitional control of movements. They are a potential control signal for brain-machine interfaces and are a therapeutic target in Parkinson’s disease. The prevailing view is that EEG beta bursts increase during stopping and immobility, but do not precede stopping. In contrast to prior work in humans and animals that used a latent and unobservable stopping time in the stop-signal task, we developed a translational animal model to align EEG with overt action stopping. We recorded 32-electrode EEG along with the angular velocity of a treadmill while head-fixed rats stopped in-progress running on a freely-rotating, non-motorized treadmill. Contrasting prior work, motor cortex beta bursts increased before stopping and not during stopping or immobility. Using information theoretic measures, we show that beta power was informative about treadmill velocity 200 msec in the future, but only during planning to stop. By introducing artificial temporal jitter to mimic the estimation of stopping time used in prior work, we show that this predictive brain-action relationship fails with even small jitter. Finally, we use a variety of machine learning methods to show that, despite EEG beta oscillations being a clear neural correlate preceding stopping, it has limited utility for real-time action decoding. Our work suggests a new conceptual model for neural control of action stopping.

## Introduction

Cortical EEG beta band (15-30 Hz) oscillation power is modulated during volitional movement. Over 75 years ago, seminal field potential recordings from the cortical surface in awake humans undergoing neurosurgery showed beta power was higher during immobility and decreased prior to and during movement^1^. Beta oscillations may maintain muscle contraction (e.g., clinching a fist or holding a posture^2^) and hence promote the “status quo*”*^3^. Externally driving beta oscillations using transcranial alternating current stimulation of human pre-supplementary motor area (pre-SMA) counteracts movement by reducing movement velocity and force^4,5^. Intrinsically generated beta bursts in human subthalamic nucleus (part of the cortico-subcortical network for volitional action control) are associated with reduced velocity of upcoming movements and slowed reaction times^6^. Both lines of evidence indicate that beta bursts have a modulatory contribution that promotes immobility and counteracts movement. While beta oscillations need to be suppressed before moving, the corollary that beta oscillations need to increase before stopping has not been clearly established. This constrains understanding how beta oscillations are a biomarker in Parkinson’s Disease and their potential as a control signal for brain-machine interfaces (BMIs).

Beta oscillations and action stopping have been studied using the stop-signal task in humans and animals. The stop-signal task dominates the field with citations approaching 10,000 per year (see Appendix 1 of ^7^). It is the chosen task for studying response inhibition in international multi-laboratory human cognitive test batteries^8^. In a study using this task, human motor cortex beta oscillation power did not change before or after stopping although pre-SMA beta oscillation power did change after stopping^9^. Beta oscillations also increase only after stopping in the subthalamic nucleus in humans performing the stop-signal task^10^. In contrast, other studies have shown subtle increases in beta power prior to stopping in human pre-SMA and frontal cortex, but not in primary motor cortex^11,12^. Yet, other human EEG studies have found that sensorimotor cortex beta band oscillations occur at the time of stopping or after stopping^13–16^, while translational EEG recordings from the macaque monkey showed no change in beta band activity prior to or during stopping ^17^. These mixed results have led some authors to conclude that beta band activity is not “causally linked” to stopping^13^ and that such signals are not useful as a stop-signal for BMIs^17^. However, the large number of contradictory studies may be due to the nature of the stop-signal task.

In the stop-signal task, a sensory signal instructs the subject to cancel a *planned* movement. There is no overt movement and no observable stopping time. Rather, the time at which the subject stops planning to move (i.e., the stop-signal reaction time, or SSRT) is estimated and thus an unobservable and hidden variable. The SSRT is mathematically modelled for each recording session using the distribution of reaction times across all trials of the recording session^18^. This task design precludes aligning brain activity with stopping in two ways. First, the model and the way the SSRT is calculated varies across studies^7^. Moreover, the results of this calculation depend on the delay between the stimulus and the stop-signal, which also varies across studies^19^. Second and critically, while the time of action stopping (and any preceding neural correlate) surely varies from trial-to-trial, the SSRT is a single time point for the entire recording session. Indeed, using electromyography from response-related muscles to estimate the covert stopping of an in-preparation movement has implied that the SSRT might be overestimated^12,20^. Testing whether beta band oscillations need to be increased before stopping – and are thus useful as a BMI stop signal – requires an overt behavioral measure of stopping an in-progress action.

Here, we present a behavioral task paradigm that permits precision alignment of beta band power with an overt, measurable stopping time. In this paradigm, rats choose to stop in-progress running and return to immobility. The peak velocity of a non-motorized treadmill reveals the time at which the rat initiates stopping. We recorded cortex-wide, 32-electrode EEG. In contrast with prior studies using the stop-signal task, we observed a single beta cycle burst at a highly consistent time prior to stopping, but not during stopping or immobility. We used information theory-based methods and machine learning to test the degree to which an overt stop time can be decoded from motor cortex EEG beta band power and show that beta oscillations are predictive of stopping on average but do so poorly on a trial-by-trial basis. Lastly, we show that there is a potential upper limit for tolerable SSRT estimation error, after which the relationship between beta band power and subsequent stopping will no longer be apparent.

## Results

We studied the relationship between EEG beta band (15-30 Hz) oscillations and spontaneous (uninstructed) overt stopping of an in-progress action in rats. We trained rats to perform a Go/NoGo task while head-fixed on a non-motorized, low-friction treadmill. Rats were required to remain immobile for 0.5 to 2.0 sec before onset of a sensory stimulus. One stimulus instructed a Go response, which was to run past a distance threshold (**Figure 1A**). The other stimulus instructed a NoGo response, which required remaining immobile for the entire duration of the NoGo stimulus (1.5 sec). During some correct rejection (CR) trials, the rat began running but chose to return to immobility before crossing the distance threshold and then sat immobile for the remainder of the stimulus presentation (**Figure 1B**). These “volitional stops” (VS) are volitional movements that consist of a stimulus-guided action followed by a non-instructed, internal decision to cancel the Go response and return to immobility. The peak velocity of these movements provides an overt, trial-by-trial measure of action stopping. We calculated response velocity from the angular position of the treadmill (32 samples/sec) and registered 55,833 CR trials with volitional stops across 306 recording sessions from 14 male rats. We divided CR trials into three groups (T3, T2, and T1) based on tertiles of peak velocity (18,577 large, 18,513 medium and 18,743 small peak velocities). The small velocity group (T1) does not contain a VS. Instead, T1 trials included those in which the rat was balancing on the treadmill producing a small movement and those in which the rat sat immobile. All analyses used the T3 group unless otherwise stated. **Figure 1C** shows the average trial velocity for volitional stops compared to false alarm trials.

**Figure 1.**
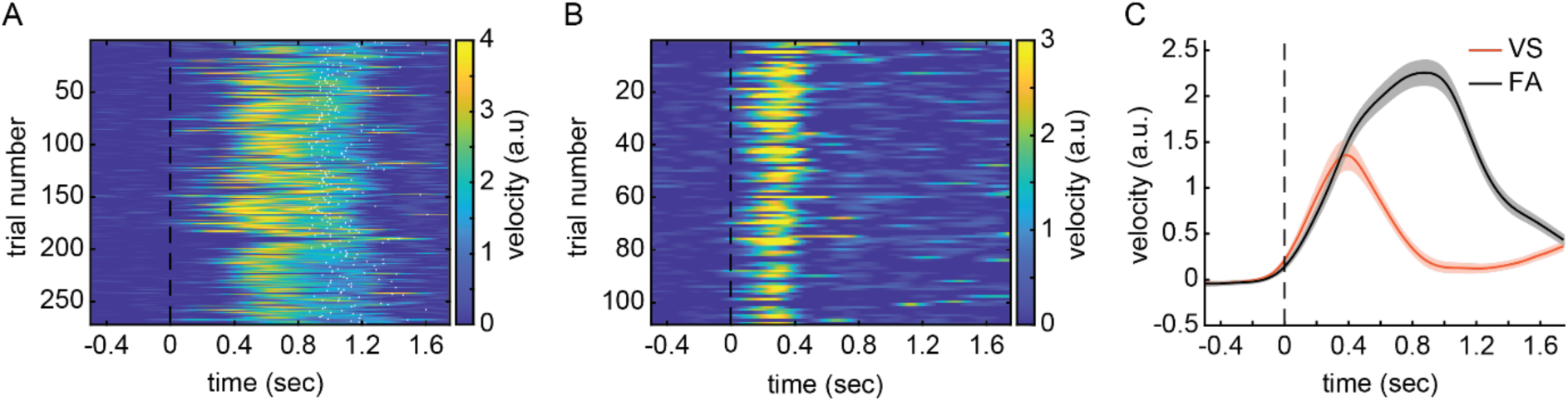
The velocity profile of volitional stopping and false alarm errors in the Go/NoGo task for the head-fixed rat on a treadmill. **(A)** NoGo stimulus-aligned velocity during false alarm trials. The data are from one session. Each row in the plot represents a trial and the white dots depicted when the response threshold (running distance) was crossed. **(B)** NoGo stimulus-aligned velocity during all CR trials in one example session. Each row in the plot is a trial. There are no white dots (as in panel B) because the rats did not commit a false alarm (FA) response. Dark blue is immobility. On numerous CR trials, the rat began running but then stopped the in-progress running before crossing the distance threshold. These VSs are visible as lighter blue, green, orange, and yellow in the plot. Variability in the peak velocity across trials is apparent and could be divided into large, medium, and small peak velocities (tertiles). **(C)** The NoGo stimulus-aligned T3 VS trial velocity (orange line) and FA velocity (black line) are plotted as an average across 14 rats. The shading shows the standard error.

We assessed how beta power changes around the time of overt stopping by recording 32-electrode EEG bilaterally and across the entire rostrocaudal extent of the rat cortex (**Figure 2A**). We calculated the power spectrum in the beta band and aligned it to trial-by-trial peak velocities. For each frequency at which we calculated power, we z-score normalized each trial to the power for the entire recording session. We assessed the topographical distribution of beta oscillations in a ± 200 msec window around peak velocity. We observed increased beta power over the left motor/somatomotor cortex (**Figure 2B**). Beta power was significantly lateralized (**Figure 2C**). In line with prior work showing that beta oscillation power decreases prior to movement onset^1,15,21^, we observed left motor/somatomotor cortex beta band power decreased during the 500 msec window prior to stimulus onset (**Figure 2D**). On the other hand, these motor-related beta oscillations were distinct from those evoked by feedback (reward or error signals), which were localized to frontal electrodes (**Figure 2E**) consistent with prior work^17,22^. Therefore, all subsequent analyses used beta band signals from these three electrodes and averaged results across them.

**Figure 2.**
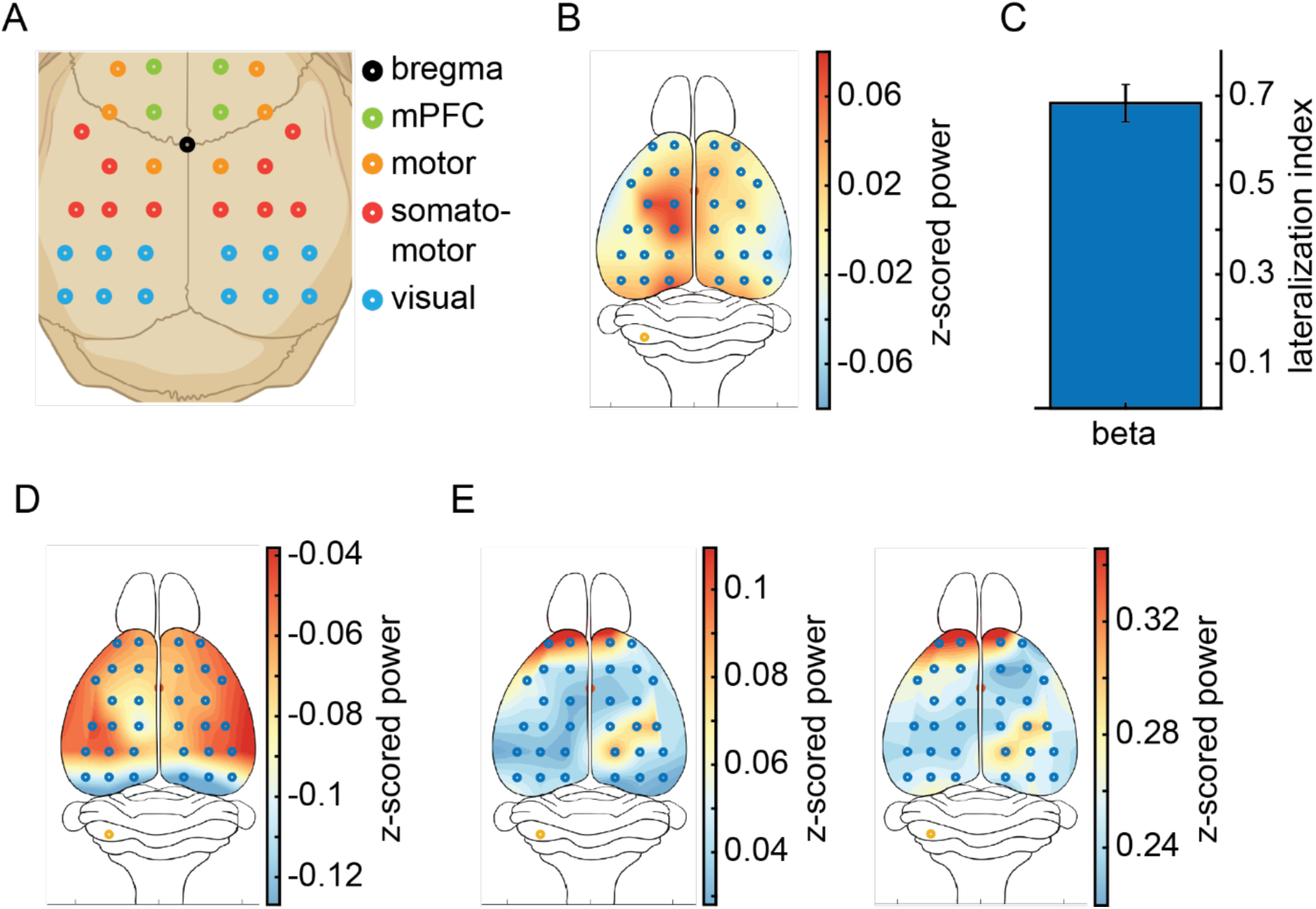
Beta power prior to stopping during VS trials has a unique cortical topography. **(A)** The schematic shows the 32-electrode skull-surface EEG array and the corresponding brain areas for each electrode. **(B)** The z-scored beta power (average, N = 14 rats) during ±200 msec window around peak velocity in T3 VS trials. **(C)** The bar plot shows the average lateralization index across 14 rats and the standard error. **(D)** The z-scored beta power 500 msec prior to stimulus onset on T3 VS trials (N = 14 rats). **(E)** The z-scored beta power in the 500 msec after reward (left plot) or auditory cue indicating that a false alarm (error) was committed (right plot). The power is averaged across 14 rats.

### Beta power transiently increases prior to stopping and positively scales with stopping larger actions

Numerous studies using the stop-signal task have shown a change in motor cortex beta power occurs too late to be predictive of stopping and instead that beta power increases during stopping (i.e., deceleration of movement) and subsequent immobility^9,11,14,15^. **Figure 3A** shows that sensorimotor cortex beta power transiently increased prior to peak velocity. The increase occurred during an epoch in which response velocity was still increasing and prior to the initiation of action stopping (black line, **Figure 3A**). Recent work has suggested that sensorimotor mu (9 – 11 Hz) oscillations can appear simultaneously with beta oscillations, and that beta oscillations can be a harmonic of the underlying mu band oscillation ^23^. We performed a simple and intuitive test for harmonics proposed by Schaworonkow (2023). If the beta oscillations are a harmonic of a mu oscillation, then the observed beta band peak frequency for each recording session will be an integer multiple of the mu band peak frequency. As proposed by Schaworonkow (2023), we used a scatter plot with a line showing the putative beta band peak frequency harmonics to visualize this cross-frequency band relationship. The points did not fall along the harmonics line and therefore the beta oscillation is genuine (**Figure 3B**). Our results indicate that beta power increases prior to overt action stopping.

**Figure 3.**
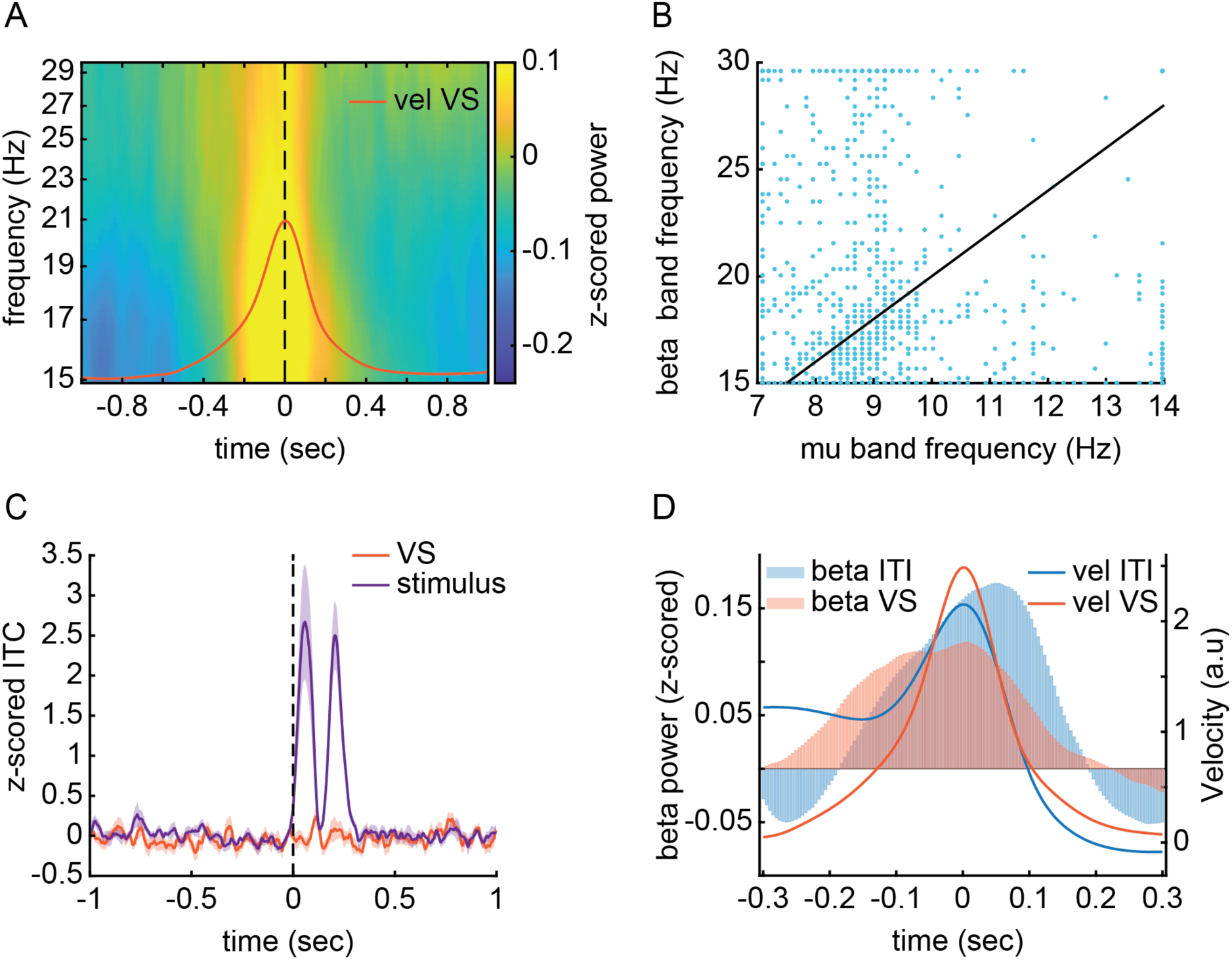
Beta power preceding VSs are a neuronal correlate of action stopping. **(A)** Averaged beta power (15-30 Hz) aligned to peak velocity on T3 VS trials. The dotted line indicates the time of the VS (i.e., peak velocity). The orange line shows the average velocity (a.u.) of the treadmill. **(B)** The beta band peak frequency is plotted against the alpha band peak frequency. Each dot is the peak frequency for a single session. The black line shows a putative harmonic relationship between the alpha band and beta band oscillations. **(C)** The lines show the z-scored ITC values averaged across 14 rats. The shading shows the standard error. The orange line is aligned to peak velocity on VS trials. The purple line is aligned with stimulus onset. **(D)** The histograms present the average (N = 14 rats) z-scored beta power aligned to peak velocity in two task epochs: VS and spontaneous task-unrelated stopping during the inter-trial interval (ITI). The orange and blue lines, respectively, show the average treadmill velocity (a.u.) for VS and stopping in-progress spontaneous running during the ITI.

However, it is a distinct possibility that beta oscillations are simply evoked by movement and phase-locked to the stop initiation event rather than occurring as an intrinsic oscillation that precedes stopping. We computed inter-trial phase coherence (ITC) to test this possibility. As a positive control, we calculated the ITC for task events in which we expected an evoked beta oscillation (i.e., after sensory stimulus onset^24–26^). Whereas we observed the expected high ITC associated with a stimulus-evoked beta oscillation, we did not observe any change in ITC around peak velocity (**Figure 3C**). Thus, the beta oscillation is an intrinsic change in brain activity prior to stopping.

It is possible that pre-stopping beta power only occurs in the context of learned stimulus-response mappings, rather than as a neuronal correlate of general action stopping. In our task, the rat has learned two stimulus-response mappings, initiates an incorrect response on CR trials, and makes a second decision to switch to the correct stimulus-response mapping before crossing the response threshold^27^. We tested whether beta power was specific to this cognitive context by measuring beta power around spontaneous running and stopping during the inter-trial interval (ITI). Rats were free to repeatedly engage in bouts of running during the ITI (at the expense of delayed stimulus onset), and they did so frequently. We calculated average beta band power aligned to peak velocity events detected during the ITI and found a similar beta power increase prior to this general stopping behavior (**Figure 3D**).

After observing that beta oscillations precede stopping, we aimed to test whether there is a predictive relationship between beta oscillations and stopping. We hypothesized that, if beta power is predictive of stopping, then there would be a relationship between the intensity of the response being stopped and the magnitude of beta power in the 200 msec prior to stopping. **Figure 4A** illustrates the power spectrum, across a wider range of frequencies (1 – 45 Hz), aligned to the time of peak velocity separately for the 3 velocity tertiles. The plots show that increasing need for stopping is associated with a larger beta band power immediately prior to stopping. We formally tested this relationship by comparing the frequency band-averaged and time-averaged (200 msec pre-peak velocity) beta power across peak velocity magnitudes (**Figure 4B**). A Bayesian independent samples one-way ANOVA indicated strong evidence for the alternative hypothesis that beta power differed with stopping magnitude (BF_10_ = 4.89×10^126^) with post-hoc t-tests indicating differences between all groups (T1 vs T2, BF_10_ = 6.55×10^42^; T1 vs T3, BF_10_ = 2.87×10^130^; T2 vs T3, BF_10_ = 4.72×10^18^). Thus, beta oscillations precede and predict subsequent action stopping.

**Figure 4.**
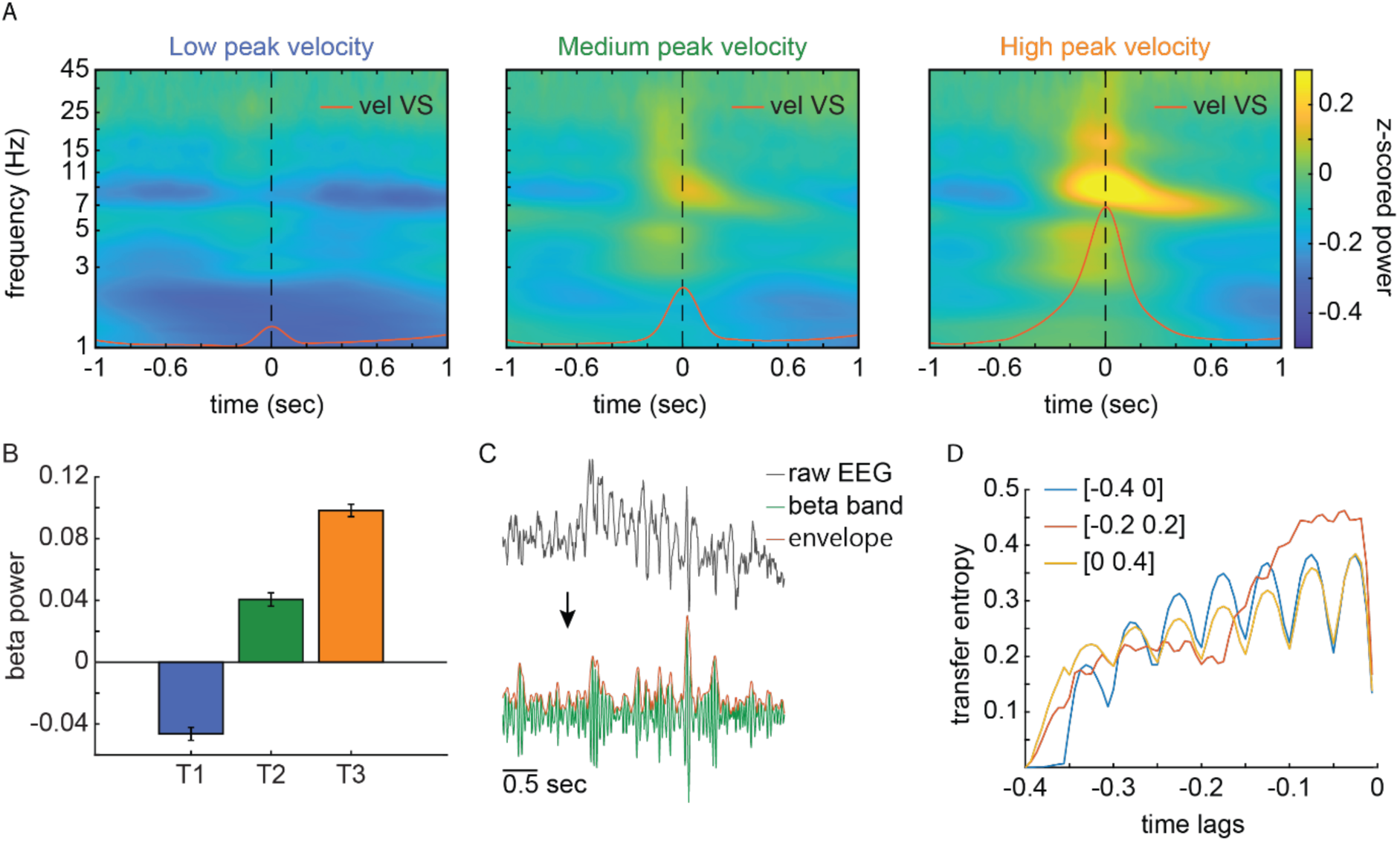
Beta oscillations precede and predict subsequent action stopping. **(A)** The power spectrum across EEG frequencies aligned to the VS. The dotted line shows the VS time (i.e., peak velocity). The orange line shows the average angular velocity (a.u.) of the treadmill in each peak velocity tertile. **(B)** Frequency band-averaged and time-averaged beta band power (z-scored to session beta band power) in the 200 msec window before the peak velocity for each velocity tertile. **(C)** An illustration of the steps used to acquire the beta band envelope. The instantaneous beta band envelope was calculated using Hilbert transformed EEG signal and then rectifying the transformed signal. The plot shows an example of raw EEG (45 Hz lowpass filtered), the beta band filtered signal, and the beta band and envelope. **(D)** Brain-to-treadmill transfer entropy in three 400 msec time windows around the peak velocity (each shown by a different line color). The different time windows consisted of distinct aspects of volitional action during CR trials: running (blue line, −0.4 to 0 sec before peak velocity), planning and execution of stopping (orange line, −0.2 to 0.2 sec around peak velocity), and stopping (yellow, 0 to 0.4 sec after peak velocity). The x-axis shows the time lag between the two signals (beta band envelope and treadmill velocity) and the lag value (*t*) should be read as the information about the treadmill at time, *t*, after the neural signal. The lines show the average across 14 rats. In the time window including both stop planning and stop initiation (orange line) only, TE increased when the time lag into the past of the beta envelope was shorter than 200 msec.

The head-fixed rat responding on a treadmill permits a brain-behavior temporal prediction analysis that is not possible when utilizing the stop-signal task. We aimed to predict the future state of treadmill velocity from the past state of beta power. We calculated brain-to-treadmill transfer entropy (TE), which is an information theory-based measure of the temporal causality between two signals^28,29^. We calculated the instantaneous beta band power envelope using Hilbert transformed EEG signal (**Figure 4C**). We then computed directional TE from the beta envelope to treadmill velocity at various time lags into the past of the beta envelope. TE increases when the past of the beta envelope yields additional information about the current state of treadmill velocity beyond the information garnered only from past velocity. Such TE increases may be interpreted as a causal influence between the signals^28,29^, in this case from the brain to the treadmill. We calculated TE in three windows of identical duration (400 msec). These windows primarily consisted of distinct aspects of volitional action during CR trials: running, planning and execution of stopping, and stopping. In the time window including both stop planning and stop initiation, TE increased when the time lag into the past of the beta envelope was shorter than 200 msec (**Figure 4D**). This indicates that the beta envelope is predictive of the state of the treadmill around 200 msec or less into the future. In contrast, in the time windows when the rats were either accelerating or executing stopping, the predictivity remained stable and lower. By using information theory to analyze the relationship between beta power and response velocity, we found that beta power is predictive of volitional stopping approximately 200 msec in the future.

### Beta oscillation bursts transiently increase in a brief window immediately prior to stopping

It has also been proposed that beta bursts occur at random times (i.e., lacking temporal regularity across trials) and that the total count of beta bursts predicts stopping^12,15^. Implicit to this conceptual model is that beta bursts are accumulated to a threshold at which point action stopping is triggered. Given that we observed a transient increase in beta power prior to overt stopping that has not been previously observed in the stop-signal task, we reassessed the beta burst accumulation theory within the context of overt stopping. We first developed a data-driven approach to find an optimal threshold for detecting beta bursts. We hypothesized that beta burst probability increases prior to peak velocity. Accordingly, setting the beta burst detection threshold too low would collect more noise, which would not be expected to change over time. On the other hand, setting the threshold too high would miss bursts and would therefore be less sensitive to change over time. **Figure 5A** shows the burst probability change over time, prior to stop initiation, for different thresholds ranging from low (the median of the beta envelope) to high (6 times the median of the beta envelope). A threshold of 2 times the median beta envelope power was the optimal criterion. Bursts were 42.4 ± 16.1 msec (SEM) duration (**Figure 5B**). This burst duration is approximately a single beta oscillation cycle (i.e., 44 msec cycle duration for the 22.5 Hz midpoint of the 15-30 Hz beta band). Approximately 50% of trials contained at least one burst in the 200 msec prior to peak velocity (**Figure 5C**). (Note that less than 100% is expected on a trial-by-trial basis when sampling a noise-containing signal). In contrast to prior findings based on the stop-signal task, a transient increase in beta bursts occurred in the 200 msec window prior to the initiation of stopping and returned to baseline immediately after stop initiation.

**Figure 5.**
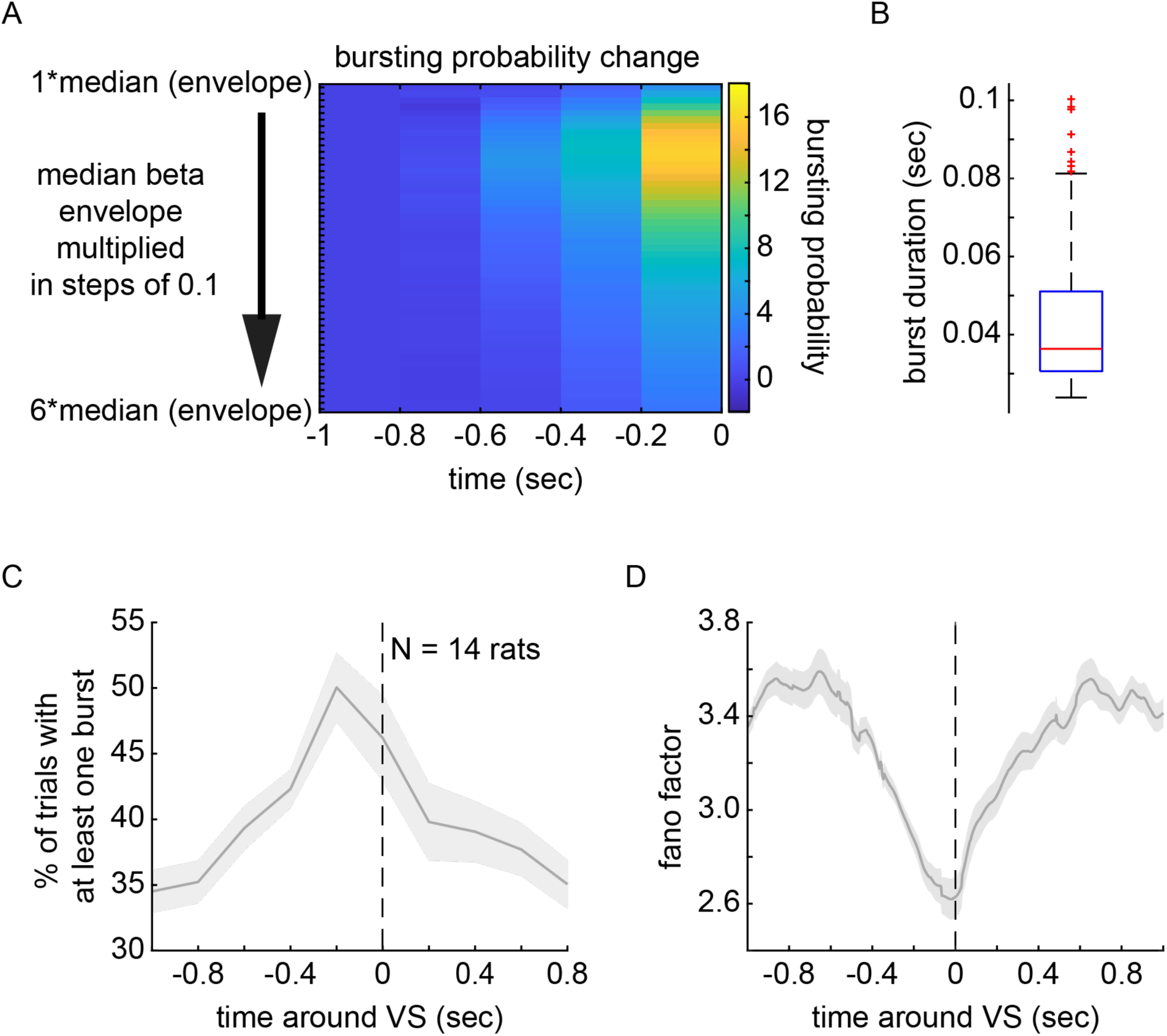
Regularly timed beta burst occurs immediately prior to stopping. **(A)** The plot shows burst probability (calculated in 200 msec bins) normalized to the first 200 msec bin during a 1 sec window prior to peak velocity. Bursts were detected using a gradually increasing threshold (steps of 0.1 times the median of the session-wide beta band power envelope) to the highest threshold tested (six times the median) at the bottom of the y-axis. **(B)** The box plot shows distribution of burst times. Outliers are marked with red crosses. The red line indicates the mean burst duration. The data set include all detected bursts in the 200 msec before VS initiation from all sessions and rats. **(C)** The percentage of VS trials with at least one burst occurring in the 200 msec window prior to the peak velocity. The dotted line shows the VS time. The line shows the average across 14 rats and the shading illustrates the standard error across rats. (**D)** The across-trial Fano Factor mean across 14 rats is plotted with the standard error shown as shading around the line. The dotted line indicates the VS time. A decrease in Fano Factor indicates a relative increase in the regularity of across-trial beta burst timing.

According to the beta burst accumulator framework, the timing of beta bursts across trials should remain random to a similar degree over time around stopping an action. We tested whether this is the case by computing the across-trial Fano Factor^30^. In contrast to this framework, we found that beta burst timing was modulated over time (**Figure 5D**). The timing of beta bursts rapidly became more regular – not more random – prior to the initiation of stopping. Importantly, beta burst rate quickly returned toward a baseline level of random timing immediately after the initiation of stopping when the rats were decelerating. This result is not in line with a conceptual model that randomly timed beta bursts are accumulated to threshold^15,31^.

### Single trial decoding of VS time from spontaneously occurring beta band power

Our results show that a single beta oscillation cycle occurs prior to volitional stopping at a highly consistent time across trials (**Figure 5B-D**). The trial-averaged power of the beta oscillation scales with the magnitude of the action to be stopped (**Figure 4A, 4B**). The ongoing, single trial beta oscillation is informative about the future state of the treadmill velocity specifically when preparing a VS (**Figure 4D**). Therefore, it is feasible that this EEG neuronal event could be used to decode the timing of an upcoming VS. We developed two decoding approaches (regression-based and classification-based) to evaluate the predictability of single trial VS time. We assessed the accuracy and robustness of a diverse range of machine learning models, encompassing both linear and non-linear methods. We pooled trials across sessions for each rat.

We first aimed to predict the exact timing, in milliseconds, of peak velocity using the beta band power envelope in the prior 400 msec. We used a 200 msec window size for the model and slid the window in increments of 10 msec (approximately one-third of a beta burst event, **Figure 5B**). To comprehensively evaluate model performance, we explored a diverse set of parametric and non-parametric regression algorithms spanning a range of complexities, including tree-based ensembles, distance-based models, and multi-layered neural networks (**Supplementary Table 1** provides a complete list of models). We elected not to include recurrent neural networks (RNNs) in our analysis because of their complexity relative to the simplicity and size of the dataset, which can cause the RNNs to overfit quickly. Instead, we substituted an RNN with a feedforward multi-layered neural network. A baseline model was included for reference (see Methods for detailed description of this model). This baseline predicted the average time-to-event from the training folds as a constant value for the test folds. We performed cross-validation and hyperparameter tuning and maintained consistency in the seeding across models (see methods section for detailed description). The evaluation metric for model selection and comparison was the average cross-validated R² score across all folds. A model predicting the mean of the dependent variable for every data point will mathematically obtain an R² of zero since it does not explain any variability beyond the average value. However, slight deviations in R² can occur when using data splits, due to small differences in the average value of the dependent variable between the training and test sets. **Figure 6A** presents the R² cross-validated scores of the best-performing model for each rat (results for all models are shown for each rat in **Supplementary Table 1**). The low R^2^ values indicate that features of the ongoing beta band power envelope alone are insufficient for precisely decoding the trial-wise VS time. While some models showed marginally better performance than the baseline—and the signal appeared more predictive for certain rats—the overall performance was poor. This suggests that the beta band power envelope lacks the necessary information for predicting VS time at the millisecond timescale. Moreover, distance-based models, such as K-Nearest Neighbors and Support Vector Machines, which rely heavily on measuring similarity between observations, struggled with the task and were never the top-performing model for any rat. This poor performance is likely due to the overlapping nature of the sequences generated by the sliding window, resulting in highly similar feature vectors despite 10 msec incremental changes in the regression value at each time step.

**Figure 6.**
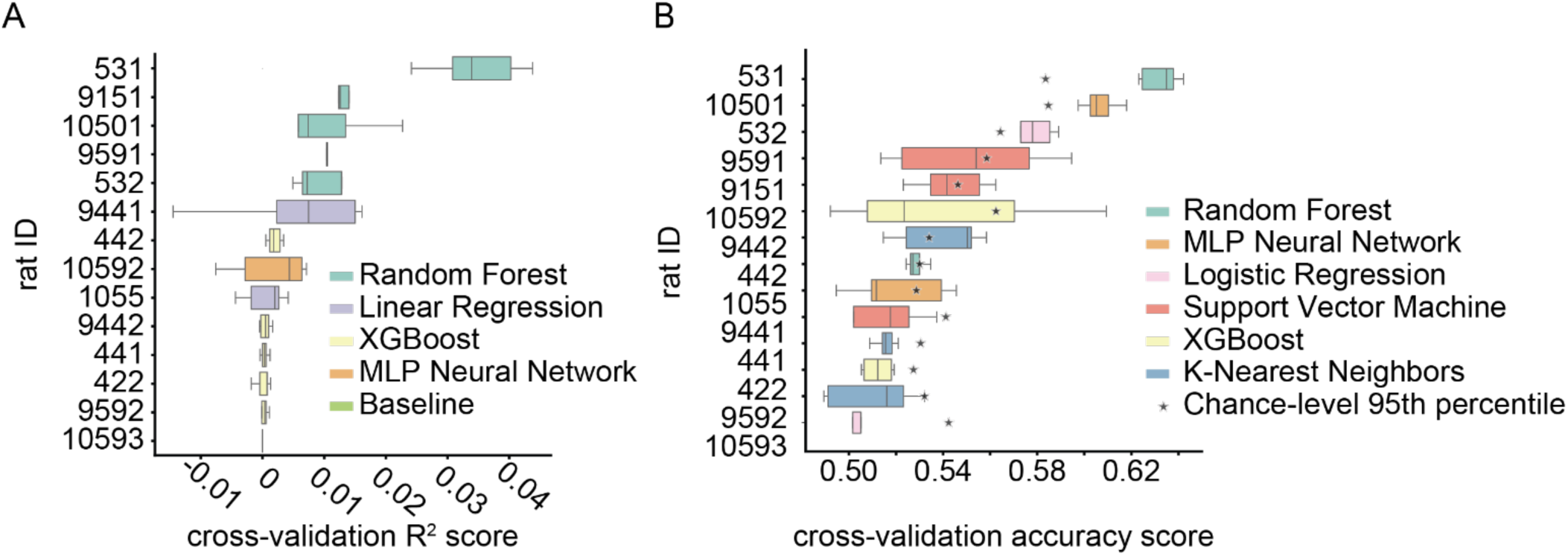
Decoding volitional stop time from ongoing EEG beta band power envelope cannot achieve high accuracy using a diverse set of parametric and non-parametric algorithms. **(A)** The distribution of cross-validated scores of the best-performing model for each rat. The regression model task was to decode the VS time on each trial. Model performance was evaluated using average R² across all folds, with the highest average R² identifying the best model. **(B)** The distribution of cross-validated accuracies for each rat’s best-performing model. The classification model task was to distinguish between the baseline and pre-VS event epoch. Overall performance was assessed by averaging accuracy across all folds, with the highest average accuracy determining the best model. For each rat, a star indicates the 95th percentile of chance-level classification accuracy. This chance level was determined by training the top-performing model using its optimized hyperparameters on shuffled training folds and then applying it to the corresponding test folds.

Therefore, we next simplified the problem and removed overlap between sequences by testing whether the beta band power envelope contained any information predictive of the VS event without focusing on precise timing. We divided the beta band power envelope, on each trial, into two time-locked segments: a baseline epoch (−400 to −200 msec before peak velocity) and a pre-VS epoch (−200 msec until peak velocity time). This approach enabled treating the problem as a binary classification task aiming to determine whether there are distinguishable beta band envelope patterns preceding peak velocity. The classification task focused on training models to discriminate between these two periods, maximizing accuracy as a performance metric. We evaluated the same range of machine learning algorithms as in the regression task (though substituting Linear Regression with Logistic Regression, see **Supplementary Table 2**). Cross-validation and hyperparameter tuning were performed similarly, creating folds accounting for individual trials and, in the case of classification, with the dataset being perfectly balanced between positive (pre-event) and negative (baseline) classes. Thus, accuracy was chosen as the primary evaluation metric since it effectively captures the models’ ability to distinguish between the two classes. The baseline model for this task was computed by shuffling class labels (see Methods section for description of this procedure). **Figure 6B** shows the cross-validated accuracy of the best-performing model for each rat (results for all models are shown for each rat in **Supplementary Table 2**). Some models achieved statistically significant accuracy levels for a sub-set of rats, indicating that the beta envelope contains some information capable of distinguishing baseline from pre-VS event sequences. However, despite this over-simplification of the problem compared to the previous regression task, these results were often statistically indistinguishable from chance. Furthermore, unlike the time-to-event results, distance-based models were often the top performers in the classification task. This suggests that when comparing distinct baseline and pre-event periods (without the overlapping data from the sliding window used in the regression task), measurable differences exist between these periods that can be captured using techniques that rely on calculating the similarity between multiple trials.

### Estimating a latent volitional stopping time can tolerate only a few hundred milliseconds of trial-wise temporal error

Here, we have shown that the beta oscillation dynamics prior to overt stopping are clearly distinguishable from prior findings that used the stop-signal task^9,11–13,15–17,24^. Inherent to the stop-signal task is that stopping occurs at a latent, unobservable time that is estimated as the identical time point across all trials in a single experiment. Naturally, this estimated stop time (the SSRT) must vary randomly with respect to the actual (and unobservable) stop time on each trial of the stop-signal task. Therefore, we next sought to introduce artificial jitter into our overt stopping times, such that we could calculate a temporal limit beyond which estimation error would occlude the beta power relationship with stopping. With this analysis, we aim to suggest what level of across-trial estimation error might be acceptable if the stop-signal task is used to seek for a relationship between beta band oscillations and stopping. We first performed 100 shuffles of each trial’s peak velocity time over a uniform interval of ±5 msec. We then calculated the average beta burst rate across these 100 shuffled peak velocity event times. After performing this procedure on each trial, we then obtained the trial averaged beta burst rate characterized by an artificial jitter of ±5 msec in the VS time. Subsequently, we repeated this procedure in 5 msec steps out to a final window of ±700 msec. This procedure assesses how much temporal jitter (from 5 msec to 700 msec) the alignment between beta bursting and VS time can handle before the relationship between the two is destroyed. We estimated the jitter duration at which this occurs by performing a one-sided t-test of the hypothesis that beta burst rate in the jittered data is less than the observed burst rate at the time point at which beta burst rate was maximal in the population (rat) average. **Figure 7** shows that randomly jittering an observable stop initiation time by more than 330 msec will likely occlude the relationship between beta burst rate and subsequent initiation of stopping. Our analysis suggests that when using a stop-signal task, if trial-by-trial variability in the SSRT exceeds 330 msec, then it will be impossible to detect beta oscillations increasing prior to stopping.

**Figure 7.**
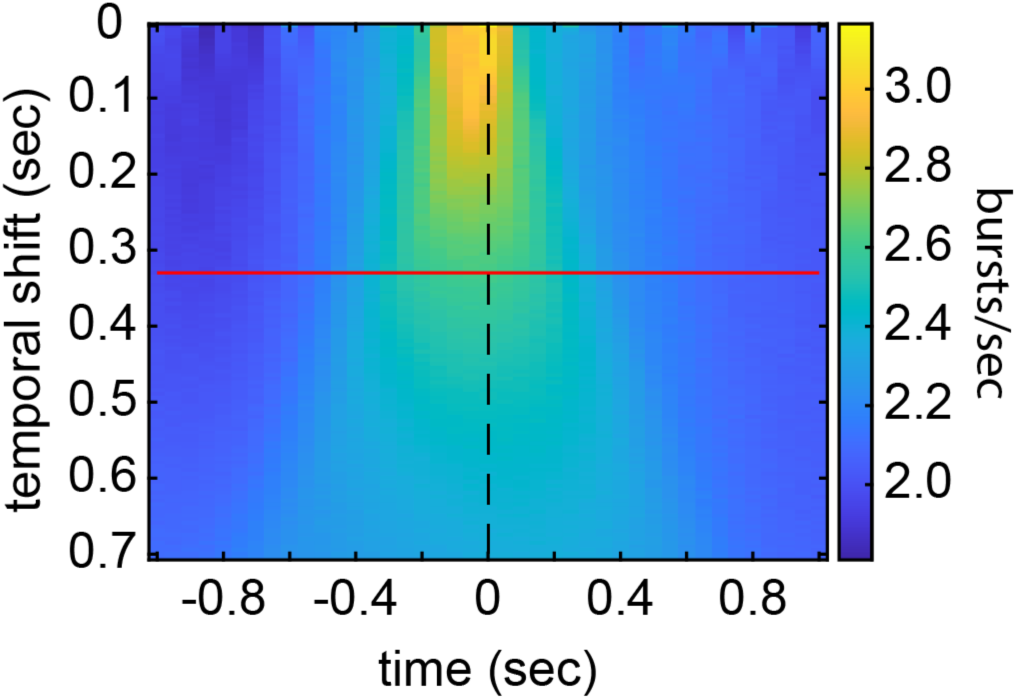
Over 330 msec error in estimating the SSRT likely occludes the relationship with beta burst and stopping. The stop initiation time was randomly jittered in 5 msec steps (y-axis). Beta burst rate was aligned to the jittered stop initiation times (time 0 on the x-axis, dotted black line). As the temporal shift grows larger (proceeding downward on the y-axis), bursts rate decreases until it is significantly lower (one-sided t-test, p<0.05) than the observed burst rate at 330 msec (red line).

## Discussion

In this study, we investigated how beta band power relates to overt stopping of an in-progress action in rats head-fixed on a non-motorized treadmill. We show a lateralized increase in beta power over the left motor/somatomotor cortex prior to stopping. Furthermore, we show that the intensity of the beta power increase is positively correlated with an increasing need to stop. The rate of beta bursts was modulated prior to stopping and this modulation occurred within a highly specific time window (200 msec before stopping). Using information theory to evaluate the predictive temporal relationship between beta power and response velocity, we found that beta power is informative about volitional stopping approximately 200 msec in the future. Information theory-based methods for inferring temporal predictions between signals were used to show that motor cortex beta power is informative about volitional stopping approximately 200 msec in the future, but uninformative about the state of the treadmill during stopping or return to immobility. Thus, a single beta oscillation precedes stopping with high trial-by-trial temporal consistency and holds predictive power over stopping. Lastly, we used synthetic data to show that, in the absence of an observable stop initiation time, estimating stopping time cannot tolerate more than 330 msec of trial-by-trial variability without losing the ability to detect a relationship between beta oscillations and subsequent stopping. This suggests a potential upper limit for tolerable SSRT estimation error, after which the relationship between beta band power and subsequent stopping will no longer be apparent.

While some studies have failed to find a relationship between beta oscillations and subsequent stopping^9,10,13–17^, those that have reported a relationship^11,12^ support two conceptual models. However, our results do not support either of them. One model suggests that randomly timed beta bursts on each trial are predictive of stopping^15^. According to this model, sustained increases in beta oscillation power only appear because bursts of one cycle (or a few cycles) of the beta oscillation occur at random times, which results in the appearance of a sustained oscillation when power is averaged across trials^31^. If randomly timed beta bursts precede stopping, then burst count must be accumulated to a threshold, which fits with long-held concepts from both spike rate neurophysiology and computational models of action selection that have suggested that the stochastic accumulation of events to a threshold triggers selection and execution of a learned action^18,32^. Instead, our data support a model in which a single beta oscillation cycle occurs at a highly consistent and non-random time immediately prior to stopping. Another model proposes that increased beta activity is associated with an *inability to change* motor programs ^24,33^. The volitional movements that we have studied may be thought of as a change from one action (running) to another action (stopping). We observed a transient increase in beta power immediately prior to stopping.

Therefore, our results suggest that beta may do the opposite of this model: beta oscillations *promote change.* We observed beta oscillations before stopping an in-progress run in two contexts: a change between two learned stimulus-response rules^27^, as well as volitional stopping during the inter-trial interval. In sum, prior models do not account for our results.

Our results not matching the current conceptual models is due to two aspects of our experiments that are unique. First, we measured an overt stopping time, which permitted precision alignment between brain activity and stopping initiation time on a trial-by-trial basis. In contrast, the SSRT is hidden from observation, and a singular across-trial estimate^7,18^. Second, we continuously measured response trajectory which allowed calculation of transfer entropy, a brain-behavior directional analysis. In the stop-signal task, response trajectory is not measured because the subject is canceling a planned movement without ever having overtly moved^4,9,11–17,20^. Presumably, the same neural correlate of stopping observed before overt stopping could be detectable in the stop-signal task, if that task had precise alignment to the latent stopping time. We tested this idea by assessing how much we could shift the stopping time before the relationship between beta oscillations and stopping is no longer detectable. This analysis resulted in an estimate of tolerable trial-by-trial error (330 msec) beyond which it would be impossible to detect the relationship. We presume that the reason prior studies using stop-signal task offer inconsistent and contradictory reports is that they may be unable to estimate the SSRT within this tolerance window.

When we aligned beta oscillations to overt stopping, our analysis of cortical topography revealed an unexpected finding. We observed lateralization of the beta power increase, which was on the same side at the population level. This might be due to engaging stopping using a preferential paw. Paw preference has been observed at an individual level in rats and there is evidence of population level preference to the right paw in a reaching task^34,35^. However, a recent meta-analysis found no population level paw preference^36^. That study proposed that, in some cases, a population level paw preference could be due to random samples of small size. On the other hand, the meta-analysis examined behavioral paw preference in (skilled) forearm reaching rather than stopping in-progress running. Moreover, in most of the studies the rat strains used were Long Evans, Sprague-Dawley or Wistar, whereas we used Lister-Hooded, which was used in only one of the studies considered in the meta-analysis. Future work can determine whether in our sample (n = 14 rats) there was a paw preference to right side, which manifested as stronger beta power in the contralateral motor cortex.

Beta band oscillations are a therapeutic target and potential biomarker in Parkinson’s disease^37–41^. Better understanding of the relation of increased beta power and stopping can be used for diagnostics and developing therapeutics. Prior work has attempted to use machine learning to decode the SSRT from ongoing EEG beta band power in humans^12^. That work reports low R^2^ values (0 to 0.03 on average) showing little-to-no ability to decode stop time from ongoing beta oscillations. However, prior attempts to decode stopping time from brain activity may have failed due to the covert nature of the SSRT. Here, we used overt stopping to test the degree to which stop time could be decoded from preceding motor cortex beta oscillation power. We used a wide variety of machine learning and pattern recognition algorithms—linear and non-linear, parametric and non-parametric—to predict time-to-stopping in milliseconds and to classify a baseline epoch from an epoch preceding imminent stopping. In contrast to prior work in which beta bursts after participants are cued to stop, we tested a more challenging scenario of predicting volitional stopping in a continuous, end-to-end fashion (the model reads the beta envelope and outputs the time-to-event using a sliding window). Despite observing a clear relationship between beta power and subsequent stopping, we found that none of a variety of models could decode trial-by-trial stop time with high accuracy. We conclude that beta oscillations are a neural correlate of stopping but are not informative enough to be utilized in a brain-machine interface. It is possible that EEG, while attractive from a translational standpoint due to non-invasive access to these signals, is not suitable for decoding due to its distance from the transmembrane currents underlying the field potential. This may be compounded by the potential for beta oscillations associated with the transmembrane currents to occur among different subsets of neurons on each instance of stopping. Single unit and local field potential recordings during the overt volitional stopping paradigm will yield an answer to this question.

Here, we have shown that a single beta oscillation cycle occurs immediately prior to stopping and that the power scales with a greater need to stop. We used an overt volitional stopping paradigm to overcome the challenges caused by the estimated SSRT and thus resolve contradictory evidence in the prior literature. We show that the two predominant models for the role of beta in volitional action control cannot account for our data. Our results suggest a novel and distinct model. Beyond a better understanding of the neural correlates of volitional actions, our results have practical implications for biomarkers, therapeutic targets, and adaptive deep brain stimulation in individuals with Parkinson’s disease^37–41^.

## Material and Methods

### Subjects

14 male Lister-Hooded rats (140-190 g body weight when implanted with electrode array and cranial chamber). Rats were pair-housed for 7 days before implantation and were thereafter single housed. Behavioral task training and experiments occurred during the rats’ active phase. The rats were housed in a reversed light-dark (07:00 lights off, 19:00 lights on) cycle. All procedures were carried out after approval by local authorities and in compliance with the German Law for the Protection of Animals in experimental research (Tierschutzversuchstierverordnung) and the European Community Guidelines for the Care and Use of Laboratory Animals (EU Directive 2010/63/EU).

### Surgery

The animal was anesthetized with isoflurane (∼1.0 – 2.0%). Heart rate was monitored throughout surgery. Buprenorphine (0.06 m/kg, s.c.), meloxicam (2.0 mg/kg, s.c.), and enrofloxacin (10.0 mg/kg, s.c.) were administered. An incision was made once the rat was no longer responsive to paw pinch. Skin and connective tissue were removed to expose the skull from the frontal bone to the neck muscle and from left to right temporal muscles. The wound margin was cauterized. The exposed bone was wiped dry and cleaned with 5% hydrogen peroxide. The bone surface was then scratched with a bone curette in a grid pattern to facilitate adhesion of the adhesive for the UV light polymerizing cement used to affix the head-fixation implant to the skull. Two component UV-curing adhesive (OptiBond, Kerr) was applied to the skull and UV cured for 30 sec at full intensity (Superlite 1300, M+W Dental). A 32-electrode polyimide electrode array (rat functional EEG, Neuronexus) was implanted aligned to bregma using an alignment mark on the array. The edges of the array were fixed to the skull using two-component dental cement (Paladur). The dental cement was used in small quantity to avoid flowing under the array between the electrodes and the skull surface. After fixation of the electrode array to the skull, a custom-made head-fixation implant was attached to the skull using UV-curing cement (Tetric EvoFlow, Ivoclar). The dental cement was bonded to the adhesive by UV curing for 60 sec at full intensity. The head-fixation implant consists of a chamber approximately the diameter of dorsal surface of the rat skull and a head-fixation post on the posterior chamber wall. A craniotomy was made over the left cerebellum, and a reference electrode (flattened 99.9% pure silver wire) was inserted through this craniotomy and laid on the surface of the dura. The craniotomy was filled with viscous, electrically conductive agar. The chamber was filled with 2-component dental cement (Paladur, Kulzer). The skin was glued to the sides of the implant using tissue glue (Histoacryl, B. Braun).

Rats recovered for 5 days after surgery. Buprenorphine (0.06 m/kg, s.c.) was administered every 12 hours for 3 days in some rats and other rats were injected with meloxicam (2.0 mg/kg, s.c.) every 24 hours for 3 days. Rehydrating and easily consumable food was provided (DietGel Recovery, ClearH2O).

### Handling and water restriction

For five days prior to surgery, the rats were handled twice a day, once in the morning and in the evening. Each session lasted at least five minutes. After five days of post-surgical recovery, access to water was restricted. During training and experiments, the rats were given 8-12 mL total water per day. Most of the water was consumed as reward during the behavioral task. The remainder of the total water volume was supplied to the rats in the cage after training. The total volume of water available daily was restricted to this level for between 5 and 14 days, while rats learned and performed stimulus discrimination experiments. After an epoch of restricted water availability, rats were provided ad libitum access to water for 24 hours.

### Head-fixation and behavioral apparatus

The rat was head-fixed on a cylindrical, non-motorized fibreglass treadmill that rotated forward or backward freely on low-friction ball bearings. The treadmill and head-fixation apparatus were inside a large Faraday cage (approximately 2 m x 2 m x 2 m) with sound proofing material. A TTL pulse-controlled pump was used to deliver 10% sucrose water via a reward port that was placed at the mouth of the rat. A computer screen (behind glass with electromagnetic shielding designed to not cause a Moire effect) in front of the rat was used to display visual stimuli covering the entire visual field of the rat. Treadmill angular position was recorded via an analog signal output from a rotary encoder (MA3-A10-125-B, US Digital) attached to the rotational axis of the cylindrical treadmill. The signal output varied between 0 V and +5 V, which mapped linearly to the rotational angle of the treadmill. The signal was sampled at 32 kHz, digitized (Neuralynx signal acquisition system), and velocity was calculated offline (in MATLAB).

### Habituation to head-fixation and behavioral task training

Habituation to head-fixation consisted of a single 20-minute session. After habituation, rats were trained to commit an instrumental response for reward. Approximately 5 uL of reward solution (10% sucrose in water) was delivered for small “shaking” or body movements on the treadmill. The threshold for triggered reward was gradually increased to train the rat to make larger body movements and eventually steps. Threshold crossings were marked with a bridging stimulus (0.1 sec duration, 500 Hz auditory tone) to aid in learning the link between movement and reward. Eventually, rats would continuously walk and receive reward. This stage required from 3 to 10 sessions (one per day). Once an animal was running and licking simultaneously (which yielded approximately 7 mL of reward solution in a session lasting 20-30 minutes), we trained the rat to make instrumental (Go) responses contingent upon the presentation of a visual stimulus.

Initially, we presented a 15 sec duration visual stimulus. The stimulus was a full field, black and white drifting grating (2.4 cycles/sec, 0.005 cycles/pixel spatial frequency, 75 deg orientation). Rats were trained to respond to the stimulus by continuously delivering reward for running during stimulus presentation. Reward delivery was triggered by crossing a threshold (a.u.) that was the same for all rats and all sessions and set at a level that was associated with bilateral locomotion. The stimulus was followed by an inter-trial interval (ITI). The ITI duration was drawn randomly from a distribution ranging from 2 to 3 sec (0.05 sec bins size).

After 2 sessions, the rats were trained to not respond prior to stimulus onset. The ITI was reduced to 1 to 2 sec, and any running that crossed a velocity threshold (manually set to capture running, same for all rats and sessions) resulted in a 0.5 sec time-out from the task and a resetting of the ITI. After one or two sessions, the rats started to suppress running during the ITI. Once this was achieved, we reduced stimulus duration in small steps (10 sec, 5 sec, 2.5 sec) over a few sessions. When stimulus duration is 2.5 sec, rats exhibit a vigorous and low-latency response upon stimulus onset. At this point in training, we reduce the ITI to 0.5 to 1 sec, and after a few sessions we reduce stimulus duration to 1.5 sec (i.e., a speeded reaction time task). Rats were given 600 trials per session. This typically yielded approximately 6 mL of sucrose solution during the task. Behavior was considered stable when omission rate was below 10%.

The Go/NoGo paradigm was introduced with the addition of a NoGo stimulus. The NoGo stimulus was at least 70 degrees different from the Go stimulus. Go and NoGo stimulus trials were delivered in pseudo-random order and in equal proportion. At most, two trials of the same stimulus type could occur consecutively. A Go response required crossing a distance threshold, which roughly corresponded to taking one step. A response offset window (0 to 0.75 sec after stimulus onset) was introduced to compensate for the pre-potent drive to respond. During this period, running did not count towards the distance threshold. This allowed low latency movements but forced the rat to appraise the stimulus and decide whether to respond. After the offset window, crossing the distance threshold caused the stimulus to disappear. Hits were rewarded (three 7 uL pulses). Responses to the NoGo stimulus led to an auditory error signal (0.5 sec duration, brown noise, 60 dB) and a time-out of 6 seconds prior to the next ITI. Training was complete when performance was above 85% and omission rate was less than 10%.

### Neurophysiological recordings

Wideband (0.1 Hz to 6 kHz) signals were recorded at 32 kHz (Digital Lynx SX, Neuralynx). Both neurophysiological signals and treadmill velocity data were down sampled to 200 Hz (unless stated otherwise).

### Volitional stops peak velocity rationale and division

To obtain the time of volitional stopping initiation on correct rejection trials (N = 55,833), which corresponds to the maximum velocity they achieved before stopping (peak velocity), we searched for the maximum velocity from stimulus onset time until 1.75 sec after. For every rat, the peak velocities were divided into three groups based on their tertiles: large peak velocity (T3, N = 18,577 trials), medium peak velocity (T2, N = 18,513 trials), and small peak velocity (T1, N = 18,743 trials). The small velocity/T1 group contained some trials in which the rat sat immobile without a volitional stop. All the further analysis is focused on the T3 group, unless stated otherwise.

### Power

The continuous wavelet transform (CWT) was calculated with the MATLAB function, cwt, in a 4 sec window around the volitional stop time to avoid window edge artifacts. The wavelet used was an Analytic Morlet Wavelet, with 48 voices per octave and frequency limits from 1 to 45Hz. Once CWT was computed, each trial was cut to a time window of −1 to +1 sec around the volitional stop time. Each time window was z-scored, within each frequency bin, using the mean and standard deviation of the entire session.

### Beta burst detection

We calculated the beta envelope for each session by bandpass filtering the EEG signal for the beta band (15 - 30 Hz) using a 2^nd^ order Butterworth filter, then performing a Hilbert transformation followed by rectification. We detected beta bursts using a data-driven thresholding approach (described in the results section). We calculated the probability of a burst occurring during the time window around the VS time in a −1 to +1 sec window around the VS. We calculated this probability in 200 msec bins for all trials. The bursting probability of the first bin (1 to 0.8 sec before VS) was subtracted from all the bins to assess how the bursting probability changed over time around the VS time.

### Inter-trial coherence (ITC)

We calculated ITC to measure phase-synchronization of beta oscillations across VS trials. We performed this analysis within sessions on the 3 sensorimotor electrodes with the largest beta power. Because trial number affects the ITC, we excluded sessions that had a total of VS trials less than the median of all sessions (median = 56). From the Hilbert transformed beta band oscillation in each session, we obtained the instantaneous phase angle of the oscillation. ITC was calculated as:

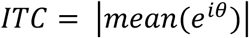

where 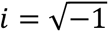 and *θ* is the phase angle. We compared the calculated ITC values against values that would be expected by chance by cutting the EEG signal at a random time point and flipping the two segments and then repeating ITC calculation. This was done 100 times to obtain a surrogate set of ITC values for each session. The mean and standard deviation of the surrogate dataset ITC values were used to calculate a z-scored ITC on each session. We also performed an identical analysis aligned to stimulus onset as a positive control in which high stimulus-evoked ITC was expected

### Fano Factor

To determine whether beta bursts occur at a consistent time or a random time across trials we calculated the Fano factor. We calculated the Fano factor for each session by multiplying the standard deviation across trials by 2 and dividing that by the mean across trials. This was done for each time point (5 msec bin size) in the time window around the VS time.

### Transfer entropy (TE)

We downsampled the two continuous signals, EEG and treadmill velocity, to 320 Hz. TE was calculated as in prior work^42^. We calculated TE in bins of 6 msec with a lag of 500 msec.

### Modeling

Machine learning algorithms were run in Python using scikit-learn. The regression-based analysis for decoding the peak velocity time used a 200 msec window, slid in 10 msec steps. Each sliding window sequence was represented as the beta envelope over time bins of N = 5 msec, resulting in 40 features per sequence. The target variable Y in this task ranged from 200 msec (for the first sequence) to 0 msec (the leading edge of the final sequence), effectively encoding the time-to-event for each window. For each rat, the sequence data was divided into five folds for cross-validation and hyperparameter tuning. To prevent data leakage from the sliding window approach, folds were created by sampling entire trials, each containing an identical number of sequences due to the time-lock applied during preprocessing. Furthermore, to maintain consistency across models, a fixed random seed was used, ensuring all models were trained on the same dataset splits. For each model, we defined a grid of hyperparameters (see code in <u><LINK></u> for details) and conducted a search for the best-performing configuration across all five folds (training on the k-1 folds and testing the model on the left-out fold, to avoid double dipping). The evaluation metric for model selection and comparison was the average cross-validated R² score across all folds. Additionally, a baseline model was included for reference. This baseline used the average time-to-event from the training folds as a constant value that we then applied for prediction over the test folds. A model predicting the mean of the dependent variable for every data point will, by definition (or the R²) obtain an R² of zero, since it does not explain any variability beyond the average value. However, slight deviations in R² can occur when using data splits, due to small differences in the average value of the dependent variable between the training and test sets.

The classification-based analysis discriminated between a baseline and a pre-VS period. Each trial yielded two labeled observations: one from the baseline period and one from the pre-event period. As in the regression task, each 200 msec sequence was divided into time bins of size N = 5 msec, resulting in 40 features that capture the beta band envelope dynamics. We evaluated the same range of machine learning algorithms as in the regression task (though substituting Linear Regression with Logistic Regression). Cross-validation and hyperparameter tuning were performed as for the regression-based analysis. However, the baseline model was calculated differently for the classification-based analysis. Chance-level performance for each model was estimated by shuffling class labels 200 times for each cross-validation train fold and running hyperparameter tuning on each shuffled dataset, effectively obtaining 200 shuffles * 5 folds = 1000 chance-level accuracies per model for the best hyperparameter configuration. This procedure removed any patterns in the data, allowing us to establish a random accuracy distribution for each algorithm. The significance of the observed performance was determined by comparing the model’s average cross-validated accuracy against the 95th percentile of the random accuracy distribution. If the observed accuracy exceeded this threshold, it indicated that the model’s performance was unlikely due to chance and suggested that the beta envelope contained information distinguishing baseline from pre-event periods.

## Acknowledgements

The authors thank Prof. Aaron Schurger for discussions about machine learning analyses and the decoding analyses performed in this paper.

---

**Supplementary Figure 1.**
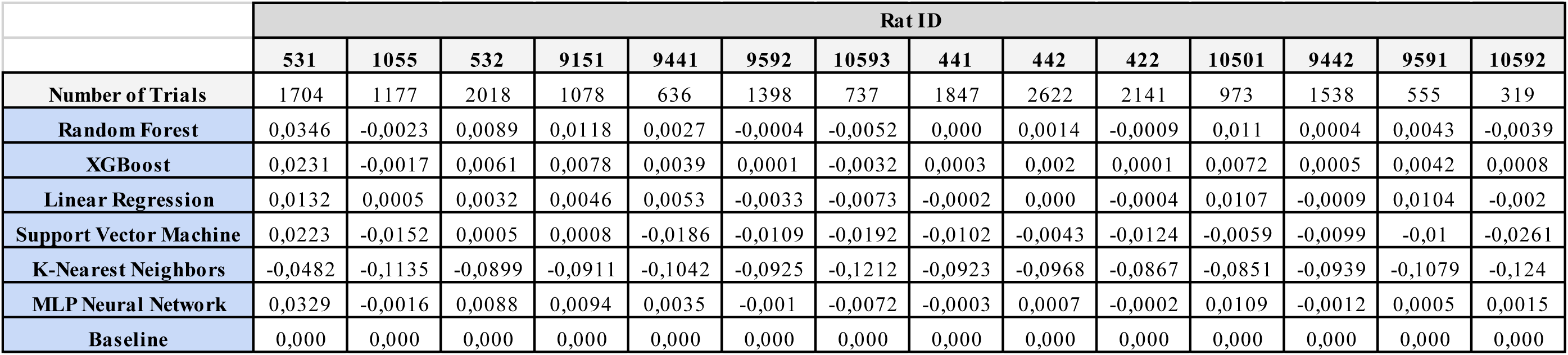
Average cross-validation R² score for time-to-event regression task.

**Supplementary Figure 2.**
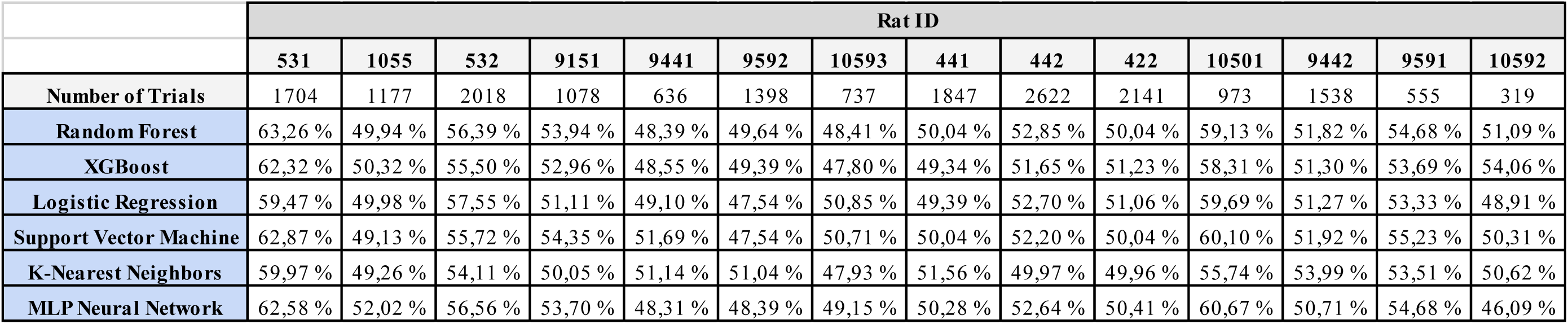
Average cross-validation accuracy score for pre-event sequence classification task.

